# Single Nucleus RNA Sequencing of Human Pancreatic Islets *In Vitro* and *In Vivo* Identifies New Gene Sets and Three β-Cell Subpopulations with Different Transcriptional Profile

**DOI:** 10.1101/2022.05.22.492974

**Authors:** Randy B. Kang, Yansui Li, Carolina Rosselot, Tuo Zhang, Mustafa Siddiq, Prashant Rajbhandari, Andrew F. Stewart, Donald K. Scott, Adolfo Garcia-Ocana, Geming Lu

**Affiliations:** Diabetes, Obesity and Metabolism Institute, and Division of Endocrinology, Diabetes and Bone Diseases, Icahn School of Medicine at Mount Sinai, New York, NY, 10029; Mindich Child Health and Development Institute, Icahn School of Medicine at Mount Sinai, New York, NY, 10029; Department of Pharmacological Sciences and Institute for Systems Biomedicine, Icahn School of Medicine at Mount Sinai, New York, NY, 10029; Genomics Resources Core Facility, Weill Cornell Medicine, New York, NY, 10065

## Abstract

Single-cell RNA sequencing (scRNA-seq) has provided valuable insights into human islet cell types and their corresponding stable gene expression profiles. However, this approach requires cell dissociation that complicates its utility *in vivo* and provides limited information on the active transcriptional status of islet cells. On the other hand, single-nucleus RNA sequencing (snRNA-seq) does not require cell dissociation and affords enhanced information from intronic sequences that can be leveraged to identify actively transcribing genes in islet cell populations. Here, we first sought to compare scRNA-seq and snRNA-seq analysis of human islets *in vitro* using exon reads or combined exon and intron reads, respectively. Datasets reveal similar human islet cell clusters using both approaches. In the snRNA-seq data, however, the top differentially expressed genes in human islet endocrine cells are not the canonical genes but a new set of non-canonical gene markers including *ZNF385D, TRPM3, LRFN2, PLUT* (β cells), *PTPRT, FAP, PDK4, LOXL4* (α cells), *LRFN5, ADARB2, ERBB4, KCNT2* (δ cells) and *CACNA2D3, THSD7A, CNTNAP5, RBFOX3* (γ cells). Notably, these markers also accurately define endocrine cell populations in human islet grafts *in vivo*. Further, by integrating the information from nuclear and cytoplasmic transcriptomes, we identify three β-cell sub-clusters: an active *INS* mRNA transcribing cluster (β1), an intermediate *INS* mRNA-transcribing cluster (β2), and a mature *INS* mRNA rich cluster (β3). These display distinct gene expression patterns representing different biological dynamic states both *in vitro* and *in vivo*. Interestingly, the *INS* mRNA rich cluster (β3) becomes the predominant sub-cluster *in vivo*. In summary, snRNA-seq analysis of human islet cells is a previously unrecognized tool that can be accurately employed for improved identification of human islet cell types and their transcriptional status *in vivo*.

## Introduction

Diabetes results from deficiency of functional pancreatic β-cells (1,2). Detailed characterization of the transcriptional programs in islet cells in health and disease will help to identify therapeutic targets to treat diabetes (3). The recent and now widely used application of single cell RNA sequencing (scRNA-seq) on human islets from healthy donors and patients with diabetes is providing a wealth of data regarding islet cell populations and their established transcriptome profile (4–10). scRNA-seq of dispersed cells from human islets or pancreas tissue represents an obvious advance over bulk RNA sequencing of whole islets or tissue and a clear improvement over bulk transcriptomic analysis of sorted islet cell subtypes, both of which require mechanical and enzymatic cell disturbances and the corresponding cellular stress. On the other hand, scRNA-seq has also disadvantages in that it also requires cell dissociation and typically focuses on exon reads which provides limited or no information on the active transcriptional status of genes since most of the mRNA analyzed is mature, stored mRNA (4).

scRNA-seq of human islets cells and human pancreas has identified genes associated with Type 1 (T1D) and Type 2 diabetes (T2D) (5–7), genes important for islet cell development and maturation (8,9), for islet dysfunction and dedifferentiation (10,11), for aging (12), and genes involved in the transdifferentiation among islet cells (13,14). These studies typically use canonical gene sets to annotate different islet cell populations, but new gene sets are continuously identified that more precisely define islet cell subtypes as they passage from development, stem cell differentiation to maturation (15). scRNA-seq has also confirmed the presence of heterogeneity among human β-cells defining several β-cell subtypes with different gene profiles (16,17). Although these are remarkable advances, it remains true that these scRNA-seq studies have been mostly performed using isolated human islets cultured *in vitro*, which may not reflect actual *in vivo* biology. Further, scRNA-seq analysis of human islet biopsies or human islet grafts in mice requires mechanical and enzymatic cell dissociation, which causes cellular stress and eliminates specific cell subpopulations sensitive to these stresses. Finally, in this *in vivo* context, scRNA-seq data sets yield very low cell numbers that may not reflect the complete universe of the normal islet cell populations (18–22).

In contrast, single nucleus RNA sequencing (snRNA-seq) uses isolated nuclei without the need for cell dissociation, can be performed on entirely intact fresh or frozen tissue, and reveals actively transcribed nascent pre-mRNAs, many of which have not yet been spliced and therefore containing introns as well as exons. Combining exonic and intronic sequences reveals important information on the transcriptional status of the cell at a moment in time (23,24). This approach should also be adaptable to analyze the transcriptome profiles and transcriptional status of human islet cells *in vivo* in previously fixed or frozen samples, as well as in human islet grafts from mice. To date, however, there are no such studies analyzing snRNA-seq in human islets *in vitro* and *in vivo* or comparing exon reads (scRNA-seq, mature mRNA) vs. exon plus intron reads (snRNA-seq, pre-mRNA) to infer the real time global transcriptional status of the cell. This is in part due to the absence of human islet cell references that include intronic data, and the absence of adequate gene sets to identify human islet cells using snRNA-seq analysis.

In this study, we addressed these latter issues by directly comparing scRNA- and snRNA-seq analysis of human islet cells to determine whether 1) this provides similar results on human islet cell populations; 2) intron plus exon reads in the snRNA-seq analysis provide additional information that further defines islet cell populations; 3) new gene sets can be defined in the snRNA-seq analysis that more accurately portray islet cell populations; 4) β-cell heterogeneity can be further defined using snRNA-seq analysis; and 5) the snRNA-seq approach can be employed for analysis of islet cell subpopulations, transcriptome profiles and transcriptional status in human islets *in vivo*. Reassuringly, the results indicate that scRNA-seq and snRNA-seq using exon or combined intron plus exon reads respectively, are interchangeable to identify human islet cell populations both *in vitro* and *in vivo*. However, the results also make the important point that at the pre-mRNA level, canonical endocrine cell genes are not the most highly expressed genes or actively transcribed in these cells, and reveal non-canonical new gene sets that accurately identify endocrine cell types in the human islet. In addition, employing combined datasets from both scRNA- and snRNA-seq analyses identifies three different β-cell subpopulations based on their *INS* gene transcriptional stage with distinct biological functions according to gene set enrichment analysis (GSEA). Overall, this study supports the use of snRNA-seq technology and pre-mRNA analysis as a tool for deciphering human islet cell populations and subpopulations and their distinct biological functions in health and disease.

## Material and Methods

### Human Islet Samples

Adult human pancreatic islets from non-diabetic donors were provided by Prodo Laboratories. The average donor age was 38±5 and 71% of them were male donors. Additional details are provided in **Supplemental Table 1**. Islets were procured in serum free medium and were cultured in non-adhesive culture plates in 5% CO_2_ at 37°C overnight before initiation of the studies.

### Human Islet Cell and Islet Nuclei Processing

Human islets [3000 islet equivalents (IEQs), 1 IEQ = 150 μm diameter islet] were collected, washed twice with PBS (Ca++/Mg++ free) and then centrifuged at 300 rpm for three min. After removing the PBS, 200 μl pre-warmed Accutase (cat# 25-058-CL, Corning) were added and the islets were incubated at 37 °C for 10 min. Then, complete RPMI medium was added to the tubes, the samples were centrifuged at 1000 rpm for three min, and the pellet washed with PBS (Ca++/Mg++ free). Half of the cells were resuspended in binding buffer (cat# 130-090-101, Miltenyi Biotec) with dead cell removal beads, incubated for 15 min at room temperature and applied onto the dead cell removal column (cat # 130-042-401, Miltenyi Biotec), which was attached to the MACS separator. Subsequently, the effluent was collected, centrifuged and resuspended with 200 μl 2% BSA and 200 U/ml RNase inhibitor in PBS. The cells were then mixed with AOPI (Cat# CS2-0106, Nexcelon Bioscience) at 1:1 ratio and the cell concentration measured with the Countess 3 Automated Cell Counter (Thermo-Fisher). The other half of the cells was homogenized with a pestle and their nuclei were isolated with the Minute™ single nucleus isolation kit for tissue/cells (Cat# SN-047, Invent Biotechnologies, INC). Briefly, cells were resuspended in 600 μl cold lysis buffer, incubated on ice for 10 min and then transferred into a filter with a collection tube. The tubes were then centrifuged at 600 x g for 5 min, the supernatants removed, and the pellets resuspended in 500μl cold washing buffer. After centrifugation at 500 x g for five min, the supernatants were removed and the nuclei pellet resuspended with 55 μl 2%BSA and 200 U/ml RNase inhibitor in PBS. Nuclei were then mixed with AOPI at 1:1 ratio and the nuclei concentration measured with the Countess 3 Automated Cell Counter. After this, nuclei samples were processed in an identical way as to the human islet cell samples.

### Human Islet Transplantation into RAG-1-/- Immunodeficient Mice

One thousand human IEQs from four different donors (**Supplemental Table 1**) were transplanted into the renal sub-capsular space of 4–5-month-old euglycemic RAGT^-/-^ mice as described previously in detail (25,26). Human islet grafts were harvested three months after transplantation, washed twice with PBS (Ca++/Mg++ free) and centrifuged at 300 rpm for three min. Nuclei were isolated as indicated above.

### Single-Cell and Single-Nucleus RNA Sequencing, Alignment and Matrix Generation

Cells and nuclei samples were prepared according to the 10X Genomics Single Cell 3’ V3.1 Reagent Kit protocol, processed with 10X Genomic Chromium Controller for partitioning and barcoding, followed by the cDNA library generation. The total cell concentration was analyzed by Countess 3, then sequenced by NovaSeq 6000 System (Illumina) at the Weill Cornell Medicine, Genomics and Epigenomics Core. FASTQ files were aligned with Cell Ranger V.6.1.1 with Single Cell 3’ V3 chemistry on the 10X Cloud’s pipeline. In the analysis, we included the intronic reads only in the snRNA-seq data with GRCh38-2020-A library. For the human islet graft samples, which were also processed identically as the snRNA-seq dataset from *in vitro* islets, we included intronic reads as well, and used the GRCh38-mm10-2020-A library to distinguish human and residual mouse genes. After the 10X h5 format file was generated, data were analyzed on the R platform with Seurat package V.4.1.1 (27).

### Quality Control, Integration and Projection

Ambient mRNA adjustment on the scRNA-seq, snRNA-seq and *in vivo* snRNA-seq data was performed using SoupX (20% contamination estimation) (28). Cells with less than a 500 genes count, less than 250 gene varieties, less than 0.8 log_10_ genes per UMI, and a greater than 20% mitochondrial gene ratio were filtered out. Then, doublets were algorithmically removed with the Doubltfinder package (20% estimation for scRNA/snRNA-seq, 10% estimation for *in vivo* snRNA-seq data) (29). Data sets were projected to publicly available Azimuth’s Human Pancreas reference (https://azimuth.hubmapconsortium.org/references/human_pancreas/) according to the script template of demo data with resulting reduction and cell-type annotation (7,27,30-35). Finally, differential expression analysis within snRNA-seq datasets for identification of new gene sets was performed.

Human *in vivo* islet data sets were projected onto our integrated scRNA/snRNA-seq data set as a reference. Since human islet grafts from harvested mouse kidneys naturally contain residual mouse cells and their mRNAs, nuclei with more than 10% of mouse genes were filtered out during quality control process.

### Unsupervised Data Analysis

After data quality control, scRNA-seq and snRNA-seq data were integrated using Seurat’s SCTransform function without allocating method parameters (30). Next, cell type identity was assigned according to the normalized gene expression level, referencing the canonical pancreatic cell type genes (27). *In vivo* snRNA-seq data were created by integration among four samples of *in vivo* snRNA-seq data and cell types were annotated by referring to both canonical markers and newly found snRNA markers from this study.

### Pathway Analysis

To define the molecular and cell function, single-cell level gene set enrichment analysis was performed using the escape package which accesses the entire Molecular Signature Database (v.7.0) (29,36-38). The whole C2 library enrichment with chemical and genetic perturbations and canonical pathways containing five databases (Biocarta, KEGG, PID, Reactome and Wikipathways) were employed. Additionally, the C5 (Gene Ontology) library was also investigated using keywords such as β-cell, pancreas, pancreatic and negative keywords such as cancer, carcinoma, anomaly or other pathologies. After enrichment scores were calculated for each single cell, they were added to the meta data for analysis and visualization.

### RNA *in situ* hybridization

RNA fluorescence *in situ* hybridization was performed on dispersed human islet cells using the RNA scope platform. Briefly, dissociated cells from human islets were plated on poly-D-lysine coated coverslips and incubated for 30 min at 37°C, 5%CO_2_. Cells were then fixed in 4% paraformaldehyde and *in situ* hybridization performed using the RNAscope^®^ Multiplex Fluorescent Reagent Kit v2, probes Hs-ZNF385D (cat#1161581) targeting intron sequences in the region 808843-810453 of NC_000003.12:22372641-21412218 and Hs-ZNF385D (cat#116501) targeting exon sequences in the region 150-1266 of NM_024697.3 which are present in *ZNF385D* pre-mRNA and mature mRNA respectively, and opal 620 (cat#FP1495001KT, Akoya Biosciences) following the instructions of the manufacturer (ACD Bio-Techne). Insulin immunolabeling was performed using anti-proinsulin/C-peptide antibody (cat# GN-ID4, DSHB), Alexa FluorTM 488 goat anti-rat IgG secondary antibody (cat#A11006, Invitrogen) and DAPI for nuclei detection.

### Statistical Analysis

Data are presented as bar graphs, violin plots and scatterplots, and show means ± SE. Statistical significance analysis was performed using t-test for comparison between groups. P < 0.05 was considered statistically significant. The simplified asterisk statistical significance annotation followed conventional criteria - 0.05, 0.01, 0.0001 and 0.0001 for increment number of asterisks.

### Study Approval

All protocols were performed with the approval of and in accordance with guidelines established by the Icahn School of Medicine at Mount Sinai Institutional Animal Care and Use Committee.

## Results

### RNA-seq Profiling of Cells and Nuclei from Adult Human Islets

**Figures 1A and 1B** depict the approach and data analysis workflow used for these studies. Islets from three healthy adult human islet donors (**Supplemental Table 1**) were used. Islet cells and nuclei from each donor were analyzed side-by-side. Cells from 3,000 islets from each of the human islet preparations were dispersed, nuclei extracted from half of the cells and the other half went through the dead cell removal process kit. After quality assessment and counting, 5,000-10,000 cells or nuclei for each sample were loaded into the 10X Genomics Chromium Controller, poly-A transcripts reversed transcribed and amplified, cDNA tagmented, and the resulting libraries sequenced to a depth of 250-500 million reads per sample (**Supplemental Table 2**). scRNA-seq and snRNA-seq data were projected onto the Azimuth human pancreas reference to determine islet cell populations and identify new gene sets as markers for these cells. In addition, scRNA-seq and snRNA-seq data were integrated and separated for the analysis of different β-cell subpopulations.

**Figure 1.**
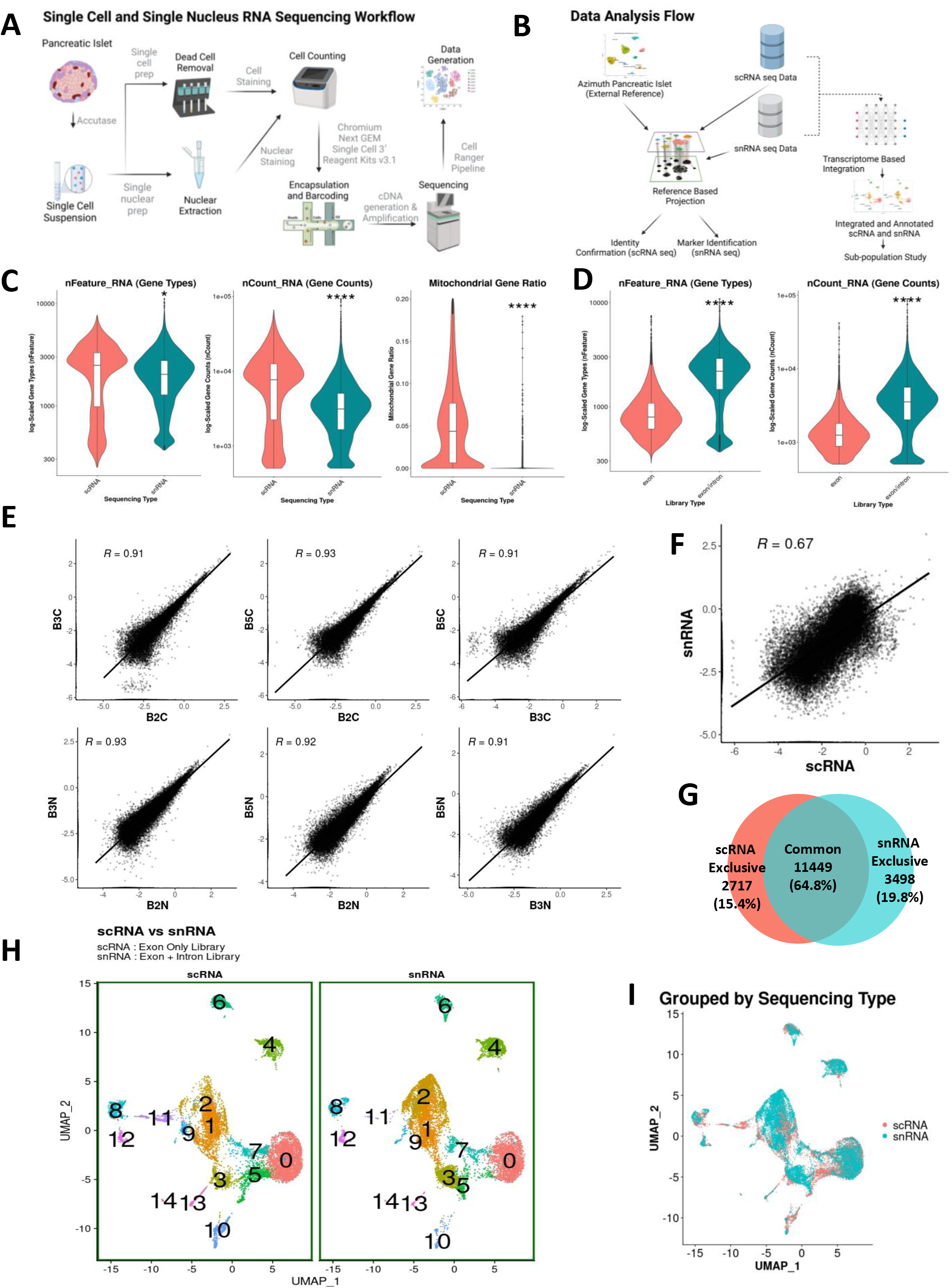
Experimental Design, Quality Assessment and Unsupervised Clustering. **A.** Human islet processing and data generation scheme. **B.** Data analysis workflow. **C.** Gene variety (nFeature), gene count (nCount) and mitochondrial ratio comparison between scRNA-seq /snRNA-seq data. **D.** Gene variety (nFeature) and gene count (nCount) comparison between snRNA data with and without intronic reads. **E.** Gene expression correlation among different human islet cell/nuclei preparations (n=3 adult human islet donors). **F.** Gene expression correlation between RNA sequencing type (scRNA /snRNA). **G.** Venn diagram of genes detected in both scRNA-seq and snRNA-seq analysis of the three human islet samples. **H.** Unsupervised clustering of scRNA-seq and snRNA-seq integrated data with Louvain resolution of 0.8. **I.** Dimensional reduction plot grouped by RNA sequencing type.

Outliers with gene counts and/or UMI count >2.5 SDs or <–2.5 SDs from the median value were excluded to eliminate cells/nuclei of inadequate quality or that represent two or more cells/nuclei. Cells and nuclei with high expression of mitochondrial genes with a cutoff of 20% were also removed. Maximum ambient RNA contamination was set up to 20% for the SoupX algorithm, and data were adjusted accordingly. Ultimately, 13,128 cells (4,395, 6,533 and 2,200 cells per human islet preparation) and 8,622 nuclei (3,033, 4,137 and 1,452 nuclei per human islet preparation) were analyzed. The number of genes and reads sequenced per cell was lower in the snRNA-seq compared to the scRNA-seq approach (**Figure 1C**) but the ratio between usable versus sequenced reads determining sequencing efficiency was similar in both methods (0.974±0.002 scRNA-seq vs. 0.966±0.002 snRNA-seq). The percentage of mitochondrial genes sequenced in the nuclei preparations were below 1% and clearly and significantly lower than the mitochondrial genes sequenced in the cell preparations (**Figure 1C**). As expected, the number of genes and reads were significantly higher when intron plus exon reads were analyzed compared with exon reads alone in snRNA-seq data (**Figure 1D**).

Based on the data quality above, we next compared scRNA-seq using exon reads with snRNA-seq using both intron and exon reads for improved gene detection and mapping as previously done (23,39). scRNA-seq analyzes both nuclear and cytoplasmic transcripts with a majority being cytoplasmic, whereas snRNA-seq profiles mostly nuclear transcripts with minimal transcripts derived from cytoplasm or rough ER during nuclei isolation (40,41). Therefore, we expected that RNA-seq reads would be different in the scRNA and snRNA sequencing profiles. In cells, 17±0.7% reads were intronic reads, while in nuclei these were 53±1.9%. On the other hand, in cells 73±1.6% reads were exonic reads in contrast to 32±0.8% in nuclei. As expected, therefore, complete or near complete linearity of gene expression correlation occurred only when nuclei were compared to nuclei or cells to cells (R=0.91-0.93) (**Figure 1E**), while this correlation was lower when cells were compared with nuclei (R=0.67) (**Figure 1F**). Indeed, the number of common genes detected by both RNA sequencing approaches was 11,449 (64.8%), while 2,717 genes (15.4%) were exclusively detected in scRNA-seq, and 3,498 genes (19.8%) were exclusively detected in snRNA-seq (**Figure 1G**). This indicates that more than 35% of the genes detected by both approaches are different for the same human islet samples predicting that the two RNA sequencing methods would reveal differences in the identity of islet cell populations. However, and contrary to this expectation, unsupervised clustering of cells and nuclei using Seurat and exonic reads (scRNA-seq) or intronic plus exonic reads (snRNA-seq) revealed clusters with similar locations and with strong overlap in their UMAPs using Seurat’s integration algorithm (**Figure 1H-I**) (42).

### Supervised Classification of Human Islet Cell Types with scRNA-seq and snRNA-seq

We next projected the scRNA-seq and snRNA-seq datasets onto a publicly available Azimuth integrated human pancreas reference that comprises six different scRNA-seq datasets generated using several different single-cell technologies using Seurat (**Figure 1B** and **Figure 2A-D**) (7,27,30-35). Projection was done with exonic reads for scRNA-seq (**Figure 2A-B**) and exonic or intronic plus exonic reads for snRNA-seq (**Figure 2C-D**). When we projected the human islet scRNA-seq data of the current study onto the reference, we found that there was a good alignment of the different human islet cell populations with a prediction score median of 1, and a mean of 0.948 (**Figure 2A-B**). Projection of the snRNA-seq data to the reference using exonic reads or intronic plus exonic reads also led to a high degree of alignment (**Figure 2C-D**) with a prediction score significantly higher with exon plus introns than with exons alone (median 0.967 vs 0.944, and mean 0.898 vs 0.861) (**Figure 2D**). This further validates the use of information from intron plus exon reads for detailed analysis of human islet cell populations using snRNA-seq. The data also indicate that human islet cell clusters from scRNA-seq and snRNA-seq share a high degree of similarity, and that snRNA-seq data containing intron plus exon reads are interchangeable with scRNA-seq data for the identification of human islet cell type clusters.

**Figure 2.**
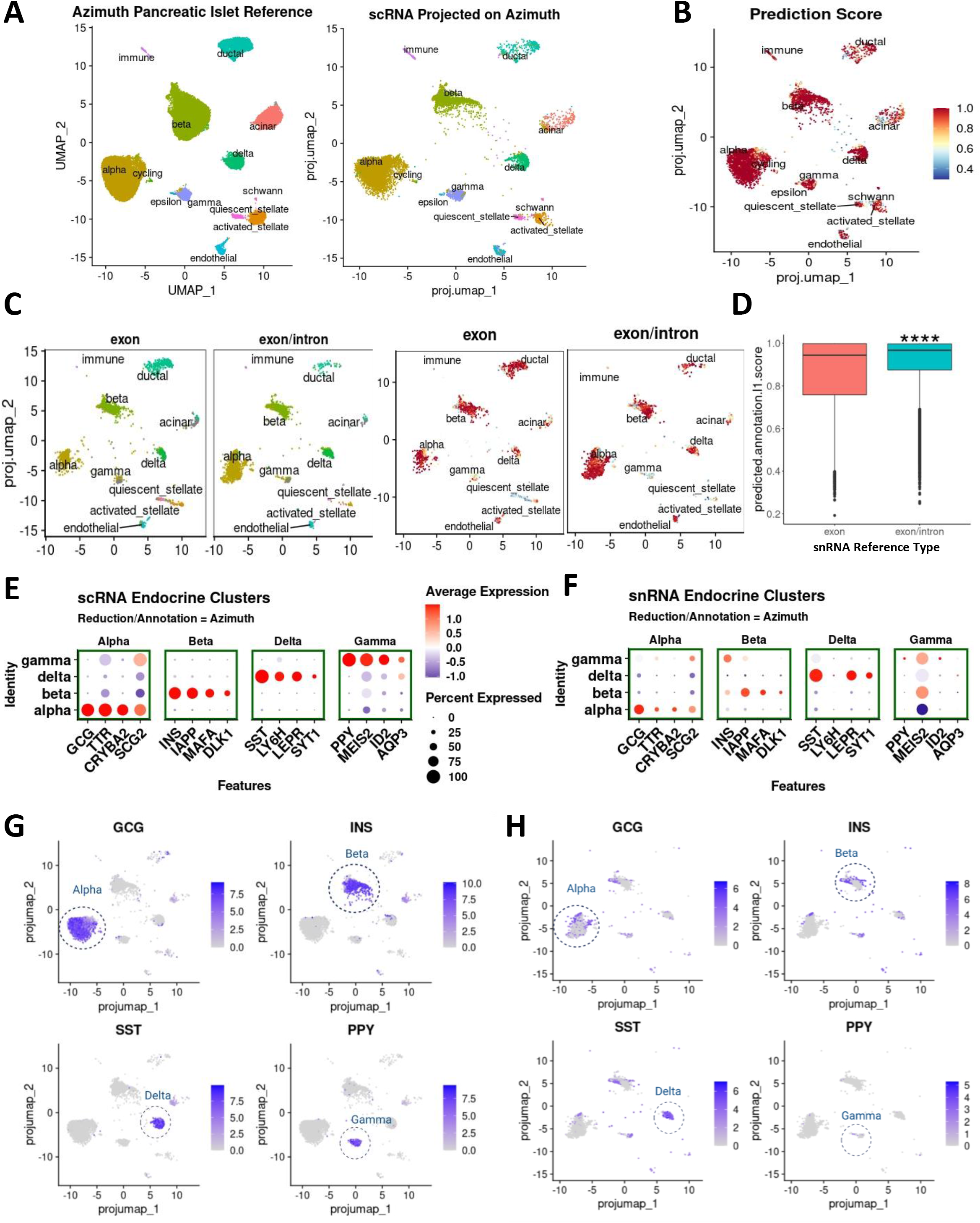
Human Islet Cell Type Identification by Projection Strategy and Mapping Score Assessment. **A.** Azimuth pancreatic islet reference v1.0.1 (left) and scRNA-seq data projected on the Azimuth reference (right). **B.** Cell type annotation prediction score displayed on dimensional reduction plot. **C.** snRNA-seq data with only exon reads or with both exon and intron reads projected on the Azimuth reference (left), cell type projection score on dimensional reduction plot (right). **D.** Prediction score in the snRNA-seq analysis with only exon reads or with both exon and intron reads in the projections in C. Statistical analysis indicates a significant (****p<0.001) higher prediction score for the snRNA-seq data projection using exon plus intron reads. **E.** Expression level and percentage of cells expressing canonical genes of Azimuth annotated endocrine cell types in the scRNA-seq data. **F** Expression level and percentage of cells expressing canonical genes of Azimuth annotated endocrine cell types in the snRNA-seq data. **G.** Dimensional reduction plot for four representative canonical endocrine genes in the scRNA-seq data. **H.** Dimensional reduction plot for four representative canonical endocrine genes in the snRNA-seq data.

Next, we tested the association of the different clusters generated from the scRNA-seq and snRNA-seq data (**Figure 2A and 2C**) with gene expression levels of canonical genes in different endocrine cell clusters (**Figure 2E-H**). A strong correlation was observed between established canonical gene cell markers (*GCG, INS, SST and PPY*) along with several other known selective markers with α, β, δ and γ cells. Thus, cell types in the UMAP were accordingly assigned in the scRNA-seq analysis (**Figure 2E and 2G**). However, the strong correlation of these canonical gene markers observed in scRNA-seq was markedly weaker in the snRNA-seq for α, β and γ cells (**Figure 2F and 2H**). For example, in beta cells, INS and IAPP are among the most highly expressed genes, whereas their expression was not impressively high in the snRNA-seq datasets. Surprisingly, this suggests that they are not the most actively transcribed genes in beta cells. Further, it suggests that annotating islet cell clusters through the expression of the canonical endocrine cell gene markers - *GCG, INS, and PPY* - is not an adequate strategy when using snRNA-seq and intron plus exon reads. Finally, it also highlights the need for the identification of new gene sets as markers for islet endocrine cells that more appropriately define them in the snRNA-seq analysis.

### Novel Gene Sets in the snRNA-seq Dataset for the Identification of Human Islet Endocrine Cell Types

We next performed differential gene expression analysis between scRNA-seq and snRNA-seq samples and investigated the biotype of the snRNA-seq enriched genes. Even with the inclusion of intronic reads, most of the genes in snRNA-seq are protein-coding genes (**Figure 3A**). Next, we annotated our snRNA-seq data with projection onto Azimuth’s human pancreas reference and tested differential gene expression for each cluster with the entire dataset (**Figure 1B**). Using this approach, differentially expressed genes in each cell cluster were identified with a p value ~0 and log_2_FC greater than 1.5. Even if the candidate genes were qualified by these criteria, we omitted genes that showed considerable expression (log_2_FC > 0.85) in other cell clusters. Thereby, we compiled a list with the top four differentially expressed genes for each cell type (**Figure 3B**). Interestingly, none of the top differentially expressed genes in scRNA-seq that define endocrine cells i.e., *INS, GCG, STS or PPY* appear in this short list of differentially expressed genes in snRNA-seq. This indicates that these canonical genes are not the most differentially expressed genes in the snRNA dataset among α, β, δ and γ cells, and therefore not ideal gene cell markers when using snRNA-seq. To confirm the reliability of these newly identified gene markers in endocrine cells, we tested them on the snRNA-seq and scRNA-seq data objects. They showed a clear and mostly exclusive pattern of expression in the corresponding cell clusters both in the snRNA-seq and scRNA-seq, but with higher expression in the snRNA-seq datasets (**Figure 3C-D**). In particular, endocrine cell markers (*PTPRT, ZNF385D, LRFN5 and CNTNAP5* for α, β, δ and γ cells, respectively) displayed a more distinctive localization pattern than their corresponding canonical single-cell clustering gene markers (*GCG, INS, SST and PPY*) for α, β, δ and γ cells, respectively in the snRNA-seq data objects (**Figure 2H and Figure 3E**). Of note, *CNTNAP5* did not display highly different expression pattern in γ cells, yet it was considered a γ cell gene marker based on high adjusted p-value (1.58×10^-5^) and log_2_FC of 1.654. Interestingly, the top differentially expressed genes in the snRNA-seq analysis in non-endocrine cells contained the canonical gene markers that define these cell types in scRNA-seq (*REG1A, CFTR, FLT1, COL1A1 and PRKG1* for acinar, ductal, endothelial, activated stellate, quiescent stellate cell, respectively). This suggests an interesting dichotomy between human endocrine and non-endocrine cells regarding the correlation between active gene transcription and steady-state transcript abundance **(Figure 3F)**.

**Figure 3.**
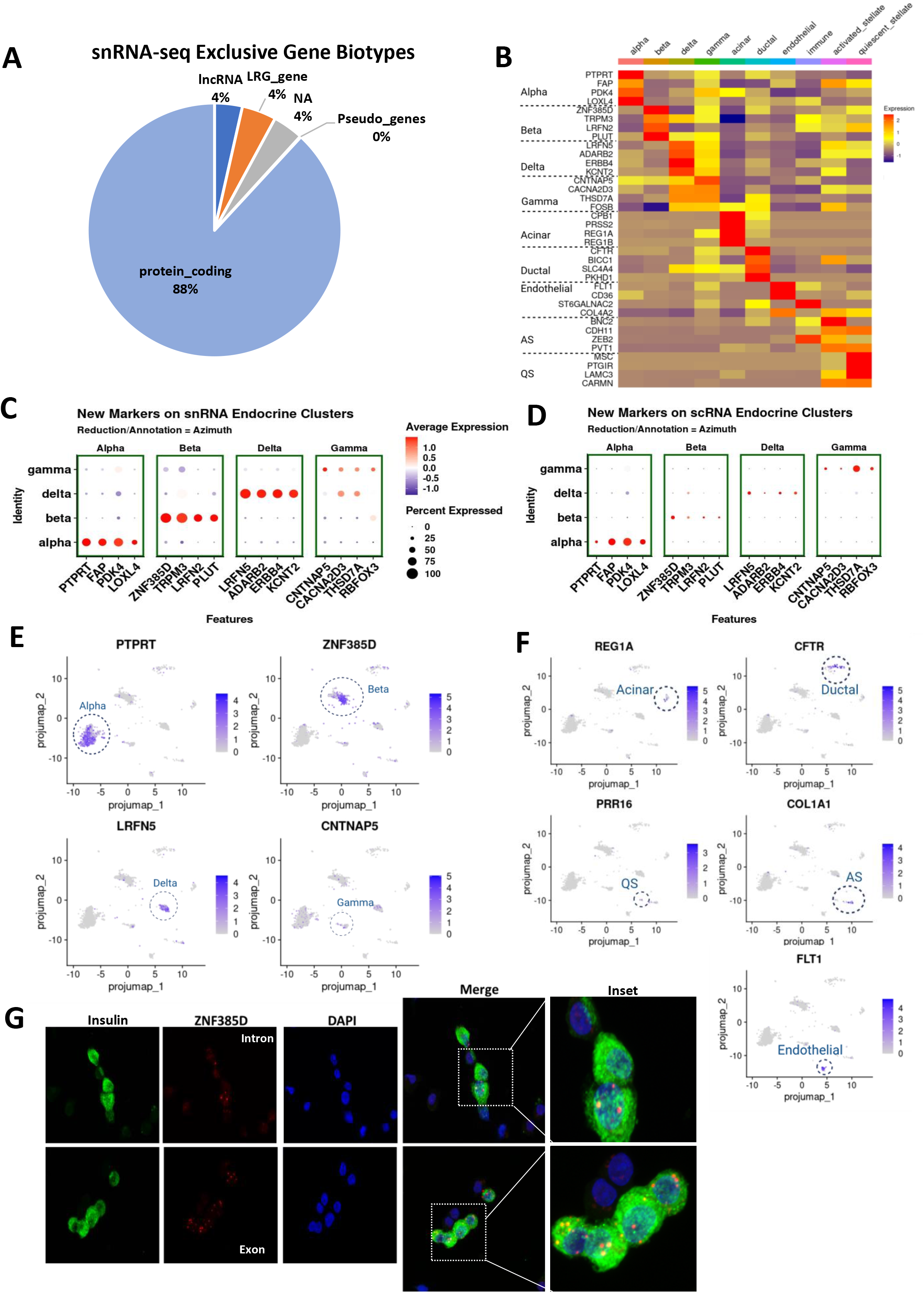
Identification of Unique snRNA-Seq Gene Markers in Human Islet Endocrine Cells. **A.** snRNA-seq identification of gene biotypes referring to AnnotationHub (54). **B.** Identification of differentially expressed genes with a p value ~0 and log2-Fold change greater than 1.5 for each cell cluster and log2FC<0.85 in the snRNA-seq data using pseudo-bulk averaged heatmap. Four top genes are presented. **C.** Dotplot projection of newly found gene markers from the snRNA-seq data on Azimuth annotated endocrine clusters. **D.** Dotplot projection of newly found gene markers in the snRNA-seq data set on Azimuth annotated endocrine clusters for scRNA data. **E.** Projection of the four top newly found gene makers for endocrine cells from the snRNA-seq data on UMAP: *PTPRT* (α cells), *ZNF385D* (β cells), *LRFN5* (δ cells), and *CNTNAP5* (γ cells). **F.** Projection of the four top gene makers for exocrine cells from the snRNA-seq data on UMAP: *REG1A* (acinar cells), *CFTR* (ductal cells), *FLT1* (endothelial cells), *COL1A1* (activated stellate cells), *PRR16* (quiescent stellate cells). **G.** Representative images of RNA fluorescence *in situ* hybridization performed in dispersed human islet cells using the RNA scope platform with probes targeting *ZNF385D* (green) intron sequences present only in pre-mRNA (top) and exon sequences present in both mature mRNA and pre-mRNA. Insulin immunofluorescence (green) was used to detect β-cells and DAPI for nuclei detection (blue).

To validate the presence of these newly identified genes from the snRNA-seq analysis as cell markers, we focused on β-cells and performed RNA scope to detect *ZNF385D* mRNA in dispersed human islet cells from healthy donors (**Supplemental Table 1**). As shown in **Figure 3G**, *ZNF385D* mRNA expression was clearly and uniquely detected in human β cells. Expression using an intronic probe was limited to the nucleus (**Figure 3G, top**) while using an exonic probe located the signal in both the cytoplasm and the nucleus (**Figure 3G, bottom**).

### Comparative scRNA-seq and snRNA-seq Analysis Identifies Three Different β-Cell Subtypes

To identify the potential existence of human β-cell subtypes within the β-cell cluster, we created a new data object integrating scRNA-seq and snRNA-seq datasets to compare insulin expression patterns (**Figure 1B and Figure 4A-C**). Combining the two datasets effectively increased the power of the analysis while providing additional information from both cytoplasmic and nuclear transcriptomes. After sub-setting the identified β-cell cluster, we generated clusters with a Louvain resolution of 0.8 (**Figure 4C**) to assign three novel β-cell sub-clusters. *INS* expression in the β-cell clusters between scRNA- and snRNA-seq data objects was different in terms of topographical location (**Figure 4D-E**). Cluster 1 displayed lower *INS* expression in scRNA-seq data but the highest in snRNA-seq data object (**Figure 4E**), suggesting that cluster 1 includes β-cells with the most active *INS* transcription since snRNA-seq analyzes mostly pre-mRNA. Next, we created a pseudo-time trajectory graph with Monocle3, assigned cluster 1 as the base (**Figure 4F**) and rearranged the order of each cluster according to the pseudo-time trajectory into β1 (active *INS* transcription), β2 and β3 cells (**Figure 4G**). Since cells in the β3 cell cluster have stable *INS* expression as inferred from the scRNA-seq (mature mRNA) β-cell cluster object, but very low *INS* expression as inferred from the snRNA-seq (pre-mRNA) β-cell cluster object, we considered these cells as the *INS*-rich cell subpopulation. Next, we looked at *ZNF385D* expression, the highest differentially expressed gene in snRNA-seq in β-cells and found minimal expression in the β-cell sub-clusters in the scRNA-seq data object but different topographical location in the snRNA-seq data object (**Figure 4H-I**) where β3 represents cells with active *ZNF385D* expression, opposite to *INS* expression (**Figure 4E**). Next, we looked at the expression of the *INS* mRNA binding protein *HNRNPA2B1,* an RNA-binding protein that regulates *INS* mRNA stability and translation (43,44), to determine whether the β2 cell cluster represents a transition stage between β1 and β3. As shown in **Figure 4J-K**, *HNRNPA2B1* expression is increased in clusters 2 and 3 compared to cluster 1, suggesting that cluster 2 may represent an “in transition” β cell type in which cells go from transcriptionally active β1 cells to β3 cells with mature stored *INS* mRNA.

**Figure 4.**
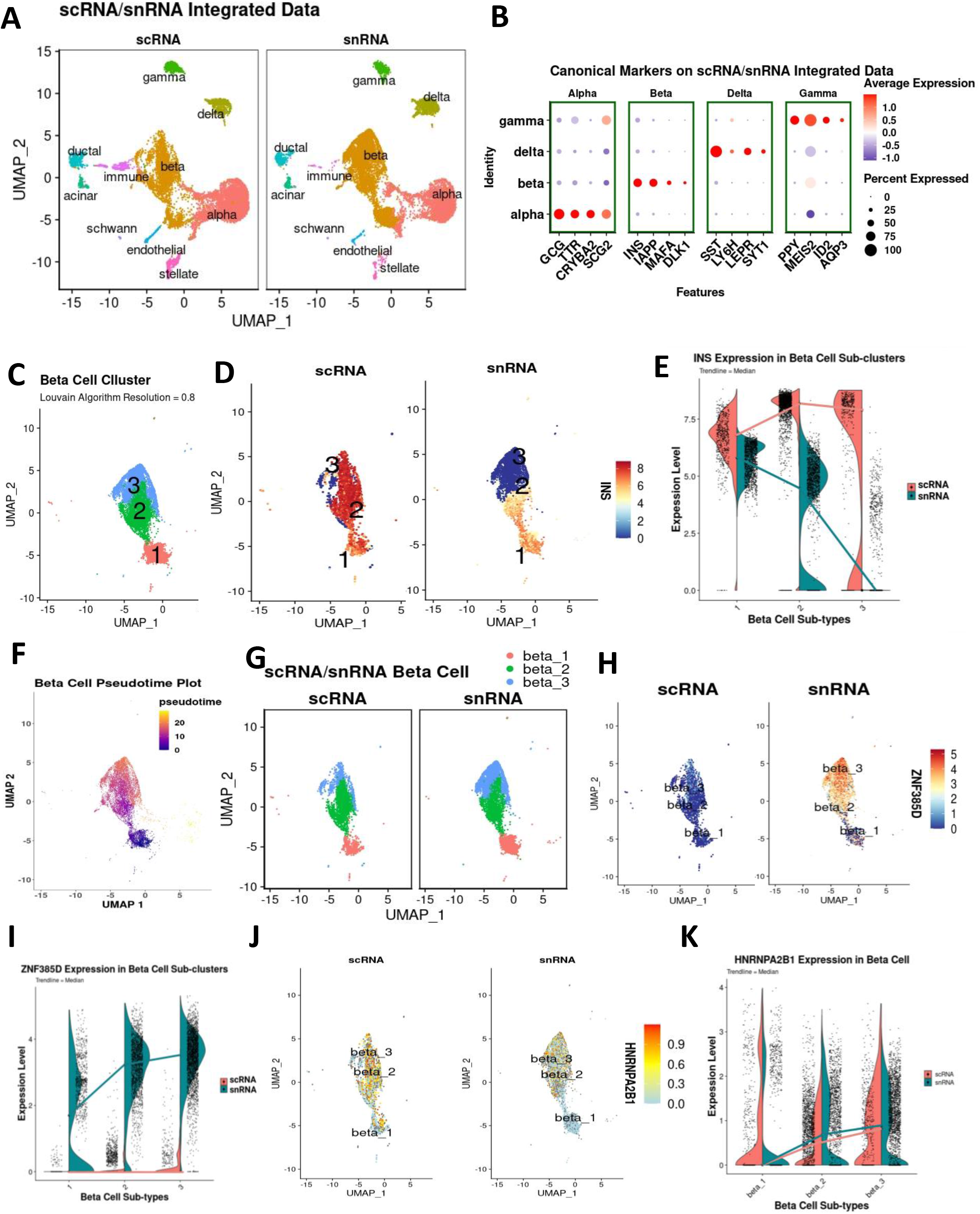
Human β-Cell Subtypes Using Unsupervised scRNA-seq and snRNA-seq Data Integration. **A.** Integrated and self-annotated clusters on UMAPs of scRNA-seq and snRNA-seq data. **B.** Canonical endocrine cell gene marker expression on scRNA-seq and snRNA-seq integrated data set. **C.** Separated β-cell cluster from integrated main data identifying three cell sub-clusters using a Louvain resolution of 0.4. **D.** *INS* expression pattern in the β-cell cluster in scRNA-seq and snRNA-seq data on UMAP. **E.** *INS* expression level on violin plots in the three β-cell subtypes in the scRNA-seq and snRNA-seq data. **F.** Monocle 3 generated pseudo-time dimensional reduction plot in the separated β-cell cluster from integrated main data using lower *INS* expression area in scRNA-seq data (cluster 1) as base. **G.** Re-annotated β-cell sub-clusters of scRNA-seq and snRNA-seq data based on pseudo-time *INS* expression. **H.** *ZNF385D* expression pattern in the β-cell sub-clusters in scRNA-seq and snRNA-seq data on UMAP. **I.** *ZNF385D* expression level on violin plots in the three β-cell subtypes in the scRNA-seq and snRNA-seq data. **J.** *HNRNPA2B1* expression pattern in the β-cell sub-clusters in scRNA-seq and snRNA-seq data on UMAP. **K.** *HNRNPA2B1* expression level on violin plots in the three β-cell subtypes in the scRNA-seq and snRNA-seq data.

### Gene Pathway Analysis in the Three Different β-Cell Subtypes

To research the potential biological differences among β1, β2 and β3, we performed GSEA of the combined datasets from scRNA- and snRNA-seq experiments (36–38). Interestingly, enrichment of genes that define the biological processes of extracellular matrix (ECM) formation, interaction and response were uniquely present in the β1 sub-cluster (**Figure 5A**). Equally, GSEA also indicated that the biological processes of intracellular vesicle budding, transport and formation are mainly present in the β2 sub-cluster (**Figure 5B**), while genes for the biological processes of insulin secretion, processing and glucose metabolism are mainly represented in the β2 and β3 sub-clusters (**Figure 5C**). Additional information was also obtained on several biological processes of importance for the development, differentiation, gene expression regulation and proliferation of the β-cell when we input the datasets from both RNA-seq approaches into the GSEA. As shown in **Figure 5D**, the genes involved in β-cell proliferation were more highly expressed in the β1 sub-cluster, the genes involved in the regulation of gene expression were prominent in the β2 sub-cluster while the genes involved in β-cell development and differentiation were more highly expressed in the β3 sub-cluster. These results clearly delineate different sets of genes for specific cell functions in the different β-cell sub-clusters emphasizing the heterogeneity of β-cells in the human islet.

**Figure 5.**
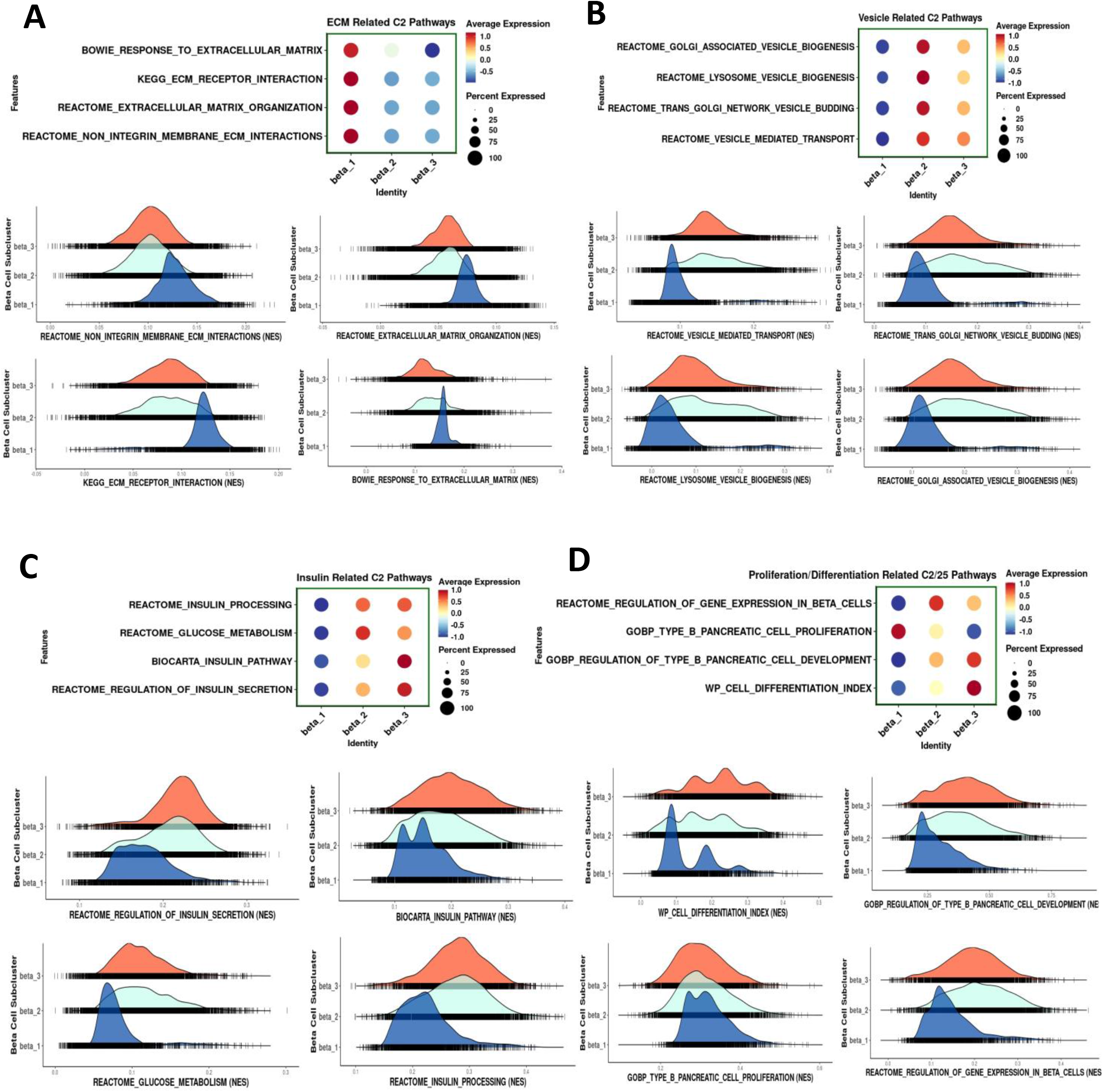
Cellular Processes Identified by Gene Set Enrichment Analysis (GSEA) of the Three β-Cell Sub-clusters β1 β2 and β3 in the scRNA-seq and snRNA-seq Integration Dataset. **A.** Dotplot depicting the expression levels and percentage of cells expressing four representative extracellular matrix (ECM) related C2 pathway genes (interaction, response, organization and integrins) and the corresponding enrichment plots below. **B.** Dotplot depicting the expression levels and percentage of cells expressing four representative vesicle-related C2 pathway genes (biogenesis, budding, transport, lysosome) and the corresponding enrichment plots below. **C.** Dotplot depicting the expression levels and percentage of cells expressing four representative insulin related C2 pathway genes (processing, secretion and glucose metabolism) and the corresponding enrichment plots below. **D.** Dotplot depicting the expression levels and percentage of cells expressing four representative proliferation/differentiation related C2 pathway genes (proliferation, development, differentiation, regulation of gene expression) and the corresponding enrichment plots below.

### Single Nucleus RNA-seq Analysis of Human Islets *In Vivo*

Next, we sought to examine the different human islet populations *in vivo* using human islet grafts transplanted in euglycemic immunosuppressed mice. For this purpose, we transplanted 1,000 human IEQs from four healthy donors into the kidney capsule of RAG1^-/-^ mice and harvested the graft three months after transplantation (**Figure 6A**). Nuclei were directly extracted from the grafts by the Minute™ single nucleus isolation kit and 5,000-10,000 nuclei per sample after quality assessment and counting were loaded into the 10X Genomics Chromium Controller, poly-A transcripts reversed transcribed and amplified, cDNA tagmented, and the resulting library sequenced to a depth of 250-500 million reads per sample (**Supplemental Table 2**). Ultimately, 7765 nuclei (1632, 1509, 713 and 3911 nuclei per human islet preparation) were analyzed. The number of gene types was 2226 ± 263, the number of gene counts was 4125.64 ± 263 and the ratio between usable versus sequenced reads determining sequencing efficiency was 0.999±0.001. The percentage of mitochondrial genes sequenced in the nuclei preparations were below 1%. snRNA-seq data were projected onto the Azimuth human pancreas reference to identify islet cell populations using canonical gene sets as well as the new gene sets described earlier (**Figure 1A and 6A**). In addition, the snRNA-seq data were projected onto the integrated scRNA-seq and snRNA-seq data from the *in vitro* studies for the analysis of different β-cell subpopulations (**Figure 6A**). Gene counts and/or UMI count outliers or cells with high expression of mitochondrial genes or with ambient RNA contamination ≥ 20% were removed. During the quality control process, we found that ≤10% mouse gene ratio was optimal for identifying different cell types *in vivo* in this islet transplant setting without considerable mouse gene influence on the clustering pattern.

**Figure 6.**
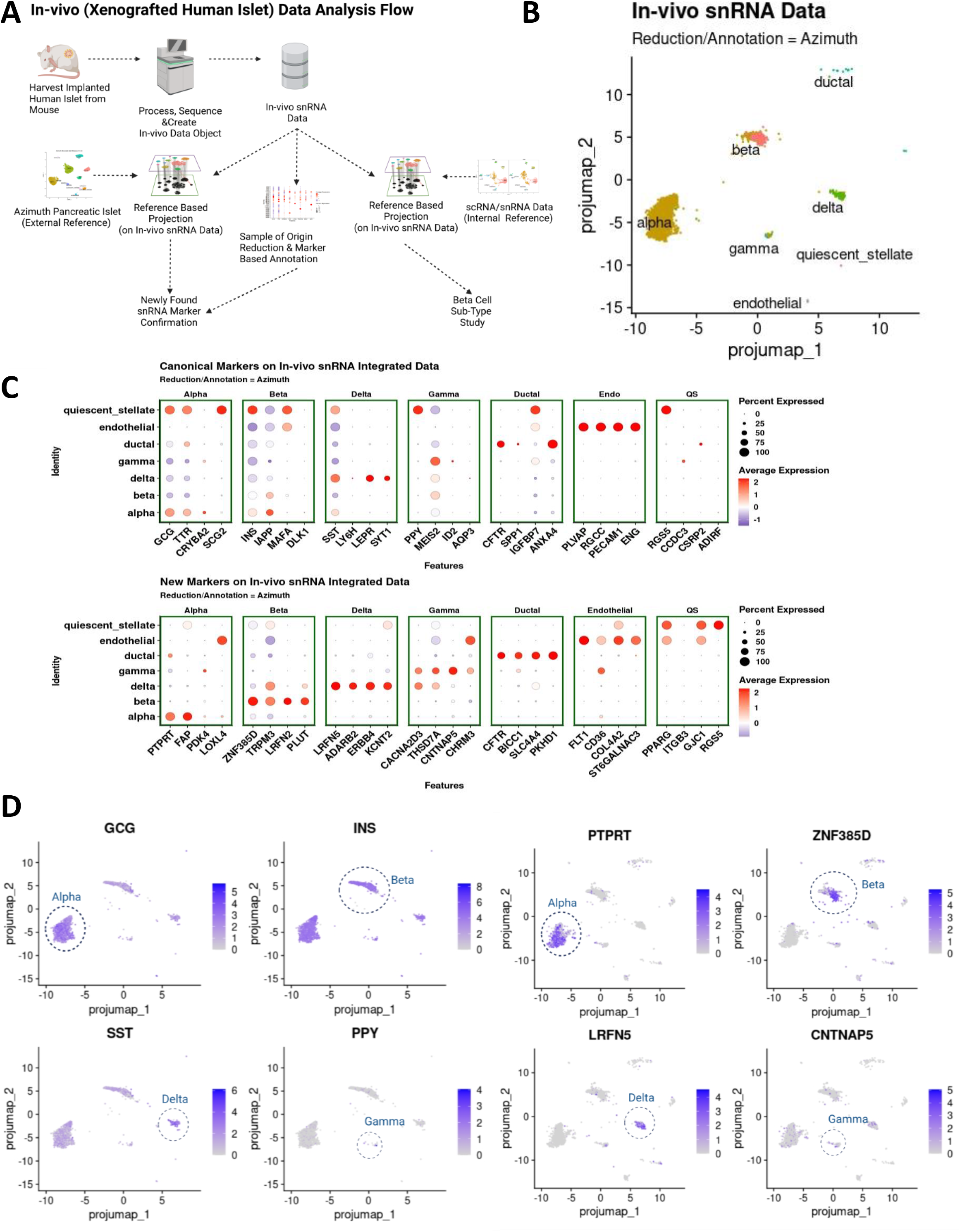
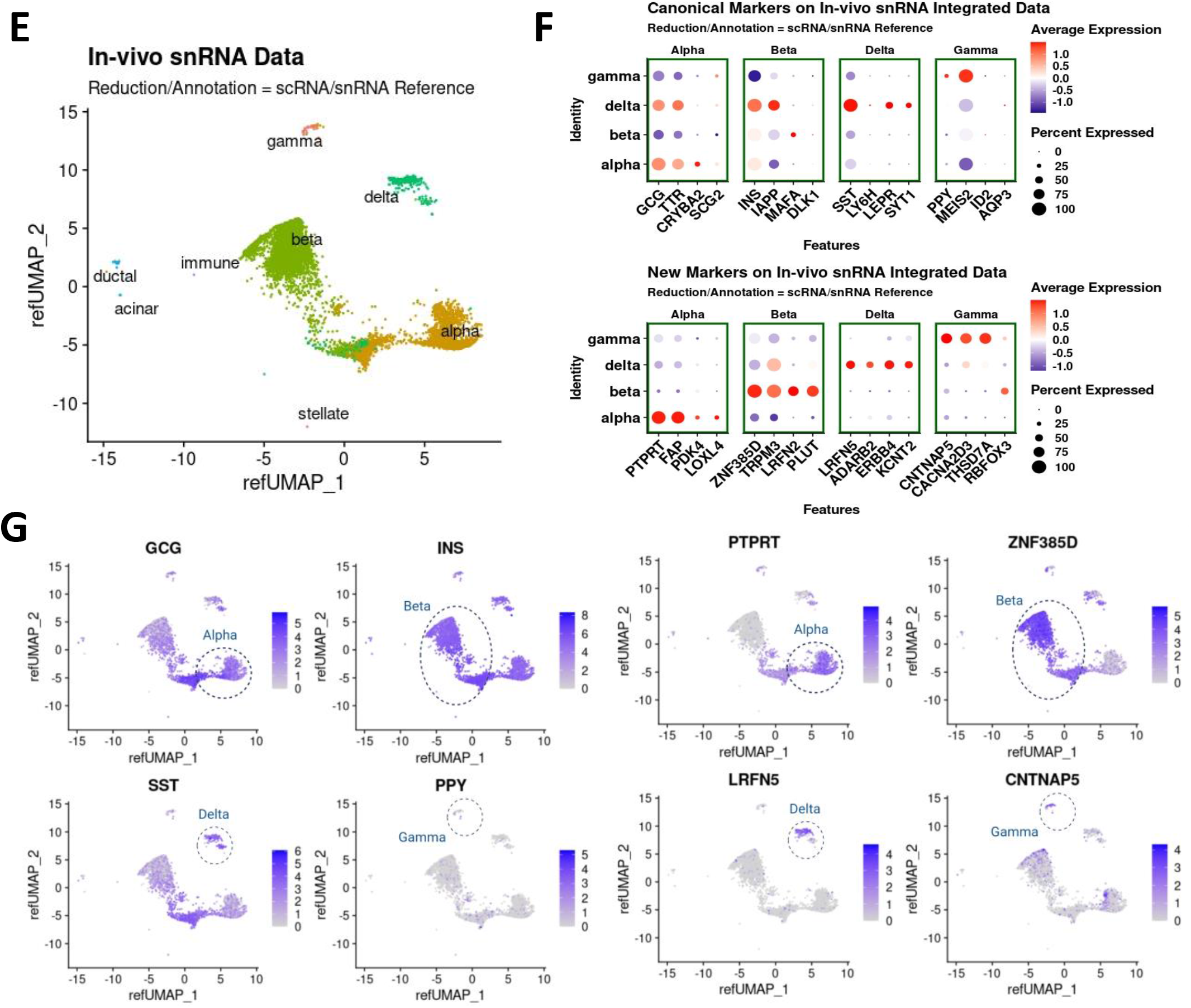
Single Nucleus RNA-seq Analysis of Human Islets *In Vivo* in Xenografts in immunosuppressed mice. **A.** Human islet grafts processing and data analysis scheme. **B.** In-vivo snRNA-seq data obtained from four different human islet xenografts done with four different human islet preparations from adult human islets from healthy donors projected on Azimuth pancreatic islet reference and cluster annotation. **C.** Dotplot depicting gene expression and percentage of cells expressing canonical gene markers (top) and newly identified gene markers for endocrine cells on Azimuth annotated in-vivo data. **D.** Projection of canonical genes and the four top newly found gene makers for endocrine cells from the *in vivo* snRNA-seq data on the dimensional reduction plot by Azimuth. **E.** *In vivo* snRNA-seq data projected on internal reference with scRNA-seq and snRNA-seq of human islets *in vitro* integrated data and annotated. **F.** Dotplot depicting the expression levels and percentage of cells expressing the canonical gene markers (top) and newly identified markers in the snRNA-seq of human islets *in vitro* (bottom) on internal reference annotated in-vivo snRNA data. **G.** Projection of canonical genes and the four top gene makers for endocrine cells found in the snRNA-seq data from *in vitro* human islets of the *in vivo* snRNA-seq data on the dimensional reduction plot by internal reference.

Using the reference-based reduction and annotation with Azimuth, we confirmed seven distinct human islet cell clusters, primarily comprised of endocrine cells (**Figure 6B**). For simplicity, we omitted clusters with less than five cells from the data set. The set of gene markers identified by the snRNA-seq analysis *in vitro* (**Figure 3C**) showed clearer patterns of specific cell expression in both the average expression-based dot plot (**Figure 6C**) and the scatterplot (**Figure 6D**) compared to the surprisingly non-specific pattern of the canonical gene markers. This specific cell cluster alignment of the newly identified gene sets from the snRNA-seq analysis *in vitro* persisted when an internal reference-based reduction and annotation (the scRNA-seq + snRNA-seq integrated datasets from the *in vitro* samples, unsupervised and marker annotated) was used (**Figure 6E-G**).

### β-Cell Subtypes in Human Islets *In Vivo*

First, we determined whether the β-cell subtypes identified in the scRNA-seq/snRNA-seq *in vitro* study were still present in human islets in the *in vivo* setting. Thus, we projected the *in vivo* snRNA-seq dataset onto the *in vitro* scRNA-/snRNA-seq dataset reference which is pre-labeled with β-cell subtypes (**Figure 6E**) and extracted the β-cell cluster (**Figure 7A**). We separated β-cell sub-clusters from the main data cluster and examined the gene expression patterns (**Figure 7A-H**). Interestingly, and in contrast to the snRNA-seq data of *in vitro* human islets, the β3 cluster in human islets *in vivo* displayed a similar level of *INS* expression compared with the β1 and β2 clusters (**Figure 7B-C**). The expression of *ZNF385D* and *HNRNPA2B1* appeared to be similar in the β-cell sub-clusters of human islets *in vivo* and *in vitro* (**Figure 7D-G**). Importantly, the proportion of cells in the β3 sub-cluster that has stable *INS* expression is significantly increased while the proportion of cells in the β2 transition cluster and the β1 with active *INS* transcription are significantly decreased in human islets *in vivo* compared with *in vitro* (**Figure 7H**). We speculate that this may represent a phenotypic or maturational transfer of cells from the lower *INS* expressing β1 and β2 groups to the more mature *INS* mRNA β3 group, presumably reflecting maturational features of the *in vivo* microenvironment as compared to the less physiological *in vitro* conditions.

**Figure 7.**
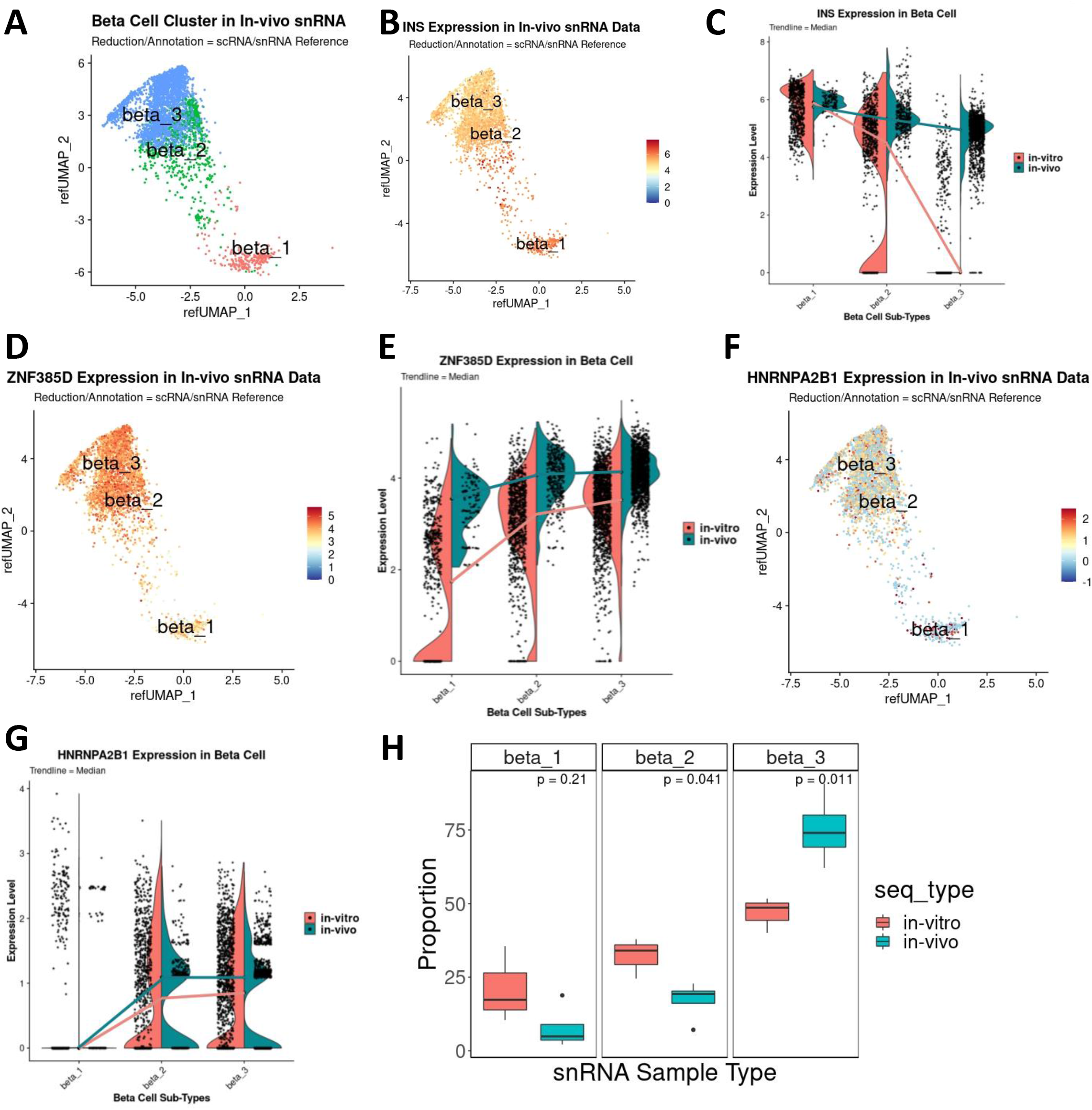
Human β-Cell Subtypes *In Vivo* from the snRNA-seq Data by Projection on the *In Vitro* Internal Reference. **A.** Separated β-cell cluster from integrated main data identifying three cell sub-clusters using a Louvain resolution of 0.4. **B.** *INS* expression pattern in the β-cell cluster in the *in vivo* snRNA-seq data on UMAP. **C.** *INS* expression level on violin plots in the three β-cell subtypes in the *in vivo* snRNA-seq data compared with the *in vitro* snRNA-seq data. Notice the difference in expression in the β3 sub-cluster. **D.** *ZNF385D* expression pattern in the β-cell sub-clusters in the *in vivo* snRNA-seq data on UMAP. **E.** *ZNF385D* expression level on violin plots in the three β-cell subtypes in the *in vivo* snRNA-seq data compared with the *in vitro* snRNA-seq data. **F.** *HNRNPA2B1* expression pattern in the β-cell sub-clusters in the *in vivo* snRNA-seq data on UMAP. **G.** *HNRNPA2B1* expression level on violin plots in the three β-cell subtypes in the *in vivo* snRNA-seq data compared with the *in vitro* snRNA-seq data. **H.** Proportion of the different β-cell subtypes *in vivo* and *in vitro*. Notice that *in vivo*, β3 cells become the majority of β-cells. Statistical analysis indicates a significant increase *in vivo* of β3 cells while β2 and β1 cell subtypes are reduced.

We also performed gene set enrichment analysis of the different β-cell subpopulations of human islets *in vivo* using the same approach as above (**Figures 5 and 8**). As observed *in vitro* (**Figure 5**), the β1 cell subtype displays higher expression levels of genes involved in ECM biological processes compared with β2 and β3 cell subtypes (**Figure 8A**). The expression of genes involved in the biological processes of intracellular vesicle budding, transport and formation, insulin secretion, processing and glucose metabolism, β-cell differentiation, development, proliferation and gene expression regulation in the β2 and β3 cell subtypes remained comparable *in vivo* and *in vitro* (**Figure 8B-D**). Notably, human β1 cells displayed lower expression of genes involved in β-cell proliferation *in vivo* than *in vitro* (**Figure 8B-D**), suggesting again perhaps that the *in vivo* microenvironment may provide cues for β-cells to favor a more functional, but less mitogenic status compared with the less physiologic *in vitro* setting.

**Figure 8.**
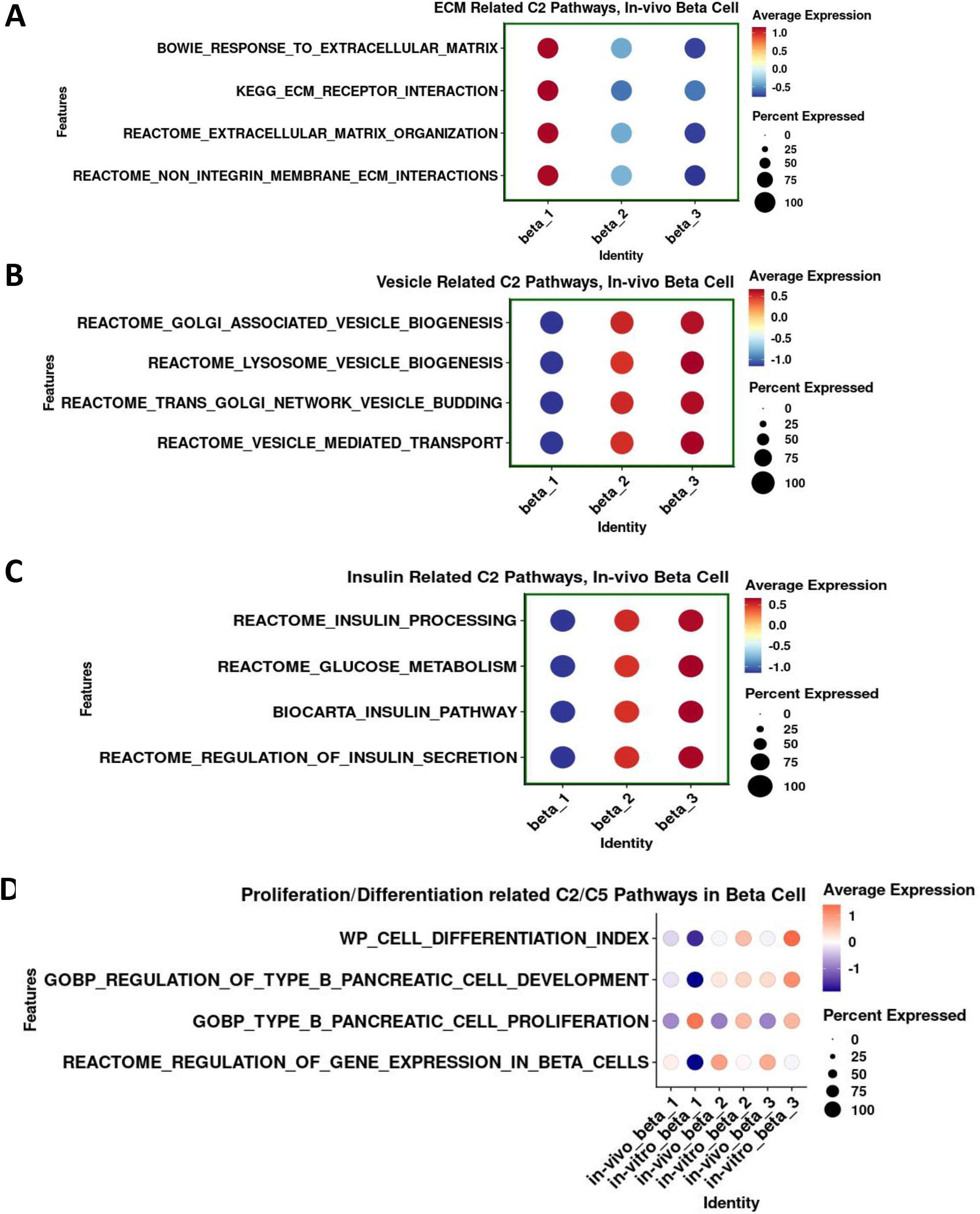
Cellular Processes Identified by GSEA of the Three β-Cell Sub-clusters β1 β2 and β3 in the *In Vivo* snRNA-seq Compared with the *In Vitro* Dataset. Dotplot depicting the expression levels and percentage of cells expressing four representative **A.** ECM related C2 pathway genes (interaction, response, organization and integrins), **B.** vesicle related C2 pathway genes (biogenesis, budding, transport, lysosome), **C.** insulin related C2 pathway genes (processing, secretion and glucose metabolism), and **D.** proliferation/differentiation related C2 pathway genes (proliferation, development, differentiation, regulation of gene expression) in the *in vivo* snRNA-seq compared with the *in vitro* dataset.

## Discussion

Approaches to accurately analyze specific human islet cell populations and their transcriptome profiles *in vivo*, immediately after harvesting the pancreas postmortem or the islet graft post-transplantation, remain an unmet need. Recently, scRNA-seq approaches have been used for such studies, but the need for tissue cell dissociation results in low cell yields, and data likely reflect cells resistant to damage and death that have been removed from their normal cell-cell contact (18-20,45,46). In many cases, only frozen or fixed human islets/pancreas samples are available.These limitiations complicate the use of scRNA-seq for *in vivo* human pancreas/islet tissue analysis. However, snRNA-seq does not require cell dissociation from the fresh tissue and can be applied to stored tissues as well (39–41). In addition, snRNA-seq can provide a wealth of information from exon and intron sequences and help to define the transcriptional status of islet cells. Here, we used snRNA-seq and analysis of intronic and exonic reads to identify human islet cell populations and their transcriptional profile and activity not only *in vitro* but *in vivo* as well. Using this approach, we defined new gene sets for annotating human islet endocrine cells by snRNA-seq and identified three human β-cell subpopulations based on their *INS* expression with different transcriptional profiles. Importantly, the proportion of these different β-cell subpopulations changes from *in vitro* to *in vivo*. These studies clealry indicate a need for further analyzing human islet cell transcriptomic profiles *in vivo*, and illustrate that the snRNA-seq technique provides a useful advance approach for this purpose.

Our goal was to establish a method that could be leveraged to annotate the human islet cell populations and their specific transcriptome profile *in vivo,* an unmet need in the diabetes field, using human islet grafts in mice. The current reference libraries to identify human pancreas cell populations and transcriptome profiles have been derived from scRNA-seq studies using mainly *in vitro* studies and exon reads since mature mRNA conforms the majority of mRNA (7,31-35). However, the majority of the reads obtained from the snRNA-seq analysis relate to intron sequences due to the pre-mRNA nature of the nuclear RNA. Based on this, we first tested whether the snRNA-seq dataset with and without intronic reads would provide similar results when projected onto the already established Azimuth scRNA-seq human pancreas reference library (7,31-35). Interestingly, inclusion of intron reads in the snRNA-seq dataset signficantly enhanced the prediciton score to the reference. Similar results have been found in other tissues as well (24), therefore we decided to use intron plus exon reads in all our analyses.

One of the surprising observations of the current study is that the expression of the main canonical genes used to define human endocrine cell populations in scRNA-seq analysis (*INS, GCG, SST and PPY*) were not the top expressed genes in the snRNA-seq studies. This indicates, first, that the most abundant genes in human islet endocrine cels are not the most actively transcribed genes in those cells. This is perhaps not surprising when considering, for example, that a large proportion of steady state cytoplasmic mRNA in β-cells is relatively stable insulin mRNA that is stored in polyribosomes ready to be translated in response to a glucose stimulus (47). Second, it indicates that new gene sets and a new reference library should be generated to use snRNA-seq analysis in human islet samples. Here, we have established this reference library and identified *ZNF385D, TRPM3, LRFN2 and PLUT* (β-cells), *PTPRT, FAP, PDK4 and LOXL4* (α cells), *LRFN5, ADARB2, ERBB4 and KCNT2* (δ-cells) and *CACNA2D3, THSD7A, CNTNAP5 and RBFOX3* (γ-cells) as new gene sets for human endocrine cells using snRNA-seq analysis. To validate at least the presence of one of these genes in human β-cells, we performed RNA scope of *ZNF385D*, the zinc finger protein 385D involved in neurocognitive development in brain (48) but whose presence and function have not been demonstrated in human β-cells thus far. Here we show for the first time that an intronic *ZNF385D* probe detects the gene in the nuclei while an exonic *ZNF385D* probe detects the gene in both the nuclei and the citoplasm of human β-cells. Future studies will decipher the role of this zinc finger protein in the human β-cell.

Another exciting result from the current studies relates to the identification of three different human β-cell subpopulations by integrating the scRNA-seq and the snRNA-seq datasets from human islets *in vitro* and using *INS* expression as the starting point in the pseudotime analysis. Heterogeneity of human β cells has been described since the 1990’s by functional analysis and more recently by scRNA-seq apporaches (16,17,49,50) but never using a combination of both scRNA-seq and snRNA-seq techniques. Since *INS* mRNA expression is higher in β1 cells in the snRNA-seq data, we postulate that β1 cells are β-cells which actively transcribe the *INS* gene. On the other hand, we postulate that since β3 cells display very low *INS* expression in snRNA-seq but very high *INS* expression level in scRNA-seq, this group of cells represent β cells which have mainy mature *INS* mRNA stored. β2 cells represent cells in a transition stage between both clusters. Interestingly, analysis of the biological processes ocurring in these β cell sub-clusters using GSEA highlightes that β1, β2 and β3 cells display different biological functions with respect to ECM, vesicle formation, insulin processing and secretion, glucose metabolism, proliferation, gene expression regulation and differentiation. This clearly places snRNA-seq analysis as an important tool to decipher heterogenitiy of defined islet cell populations.

Recently, analysis of human islet grafts by scRNA-seq has been reported with the identification of α, β, and δ cell subsets (20). However, this study used a large amount of islets (4000) transplanted in mice from where approximately only 700 cells were retrieved and analyzed, further emphasizing the difficulties of dissociating cells from human islet grafts for scRNA-seq analisys and the potential selection of cells that are resistant to mechanical and enzymatic cell disturbances with their corresponding cellular stress. More recently, snRNA-seq analysis of human islet grafts transplanted in mice has been reported (22). In this study, human islet cell types were annotated using only exon reads without using the information that the intron reads can provide to define the transcriptional status of the annotated islet cell populations. Here, we used the novel gene sets identified in the snRNA-seq analysis of human islet cells *in vitro* to annotate the human islet cell clusters in the islet grafts and their transcriptome profile. Furthermore, we were able to confirm the presence of three β-cell subpopulations *in vivo* which display the same biolgical processes described *in vitro*. Interestingly, however, we found that β3 cells are the most prominent β-cell subtype in the human islet graft. Although biological processes are maintained in the β-cell sub-clusters, it is important to note that the β-cell proliferation biological process is not present in the β1 sub-cluster *in vivo* indicating several points. First, that β-cell proliferation capacity is reduced *in vivo*, an aspect already described in several studies testing β-cell proliferation inducers for regenerative purposes which describe a lower response to these inducers *in vivo* in β-cells (25,26,51). Second, that dynamic changes occur among the different β-cell subpopulations when comparing *in vitro* and *in vivo* situations. And third, that β1 cells have higher proliferative capacity but low expression of genes involved in the insulin secretion pathway while β3 cells display the opposite phenotype highligting the dichotomy proliferation-function that occurs in β-cells (52,53). Finally, the use of snRNA-seq and β-cell subpopulations analysis *in vivo* in islet grafts opens up the future for a plethora of studies analyzing the effect of drugs and physiological and pathological situations *in vivo* in human islet cells using this genomics technique.

In summary, this study clearly supports the use of snRNA-seq analysis to define cell populations, transcriptome profiles and transcriptional status of islet cells in human islets/pancreas tissue *in vivo*. Using this approach, we have found novel gene sets that define islet cell populations in the snRNA-seq studies that can be used for islet cell identification and transcriptome profile *in vivo*. We have unraveled three β-cell subopulations with dynamic biological profiles and activities in basal conditions, that can be further interrogated *in vivo* in physiological and pathological situations related to diabetes.

## Author Contributions

R.K., G.L., P.R., A.F.S., D.K.S. and A.G.-O. conceived of the studies, oversaw the experiments and wrote the manuscript. G.L., R.K., Y.L., C.R., T.Z. and M.S. performed the studies.

## Acknowledgements

We thank the NIDDK-supported Einstein-Sinai DRC and the Human Adenovirus and Islet Core for help with the proposed studies and Prodo Labs for supplying human cadaveric islets. This work was supported by NIH grants P-30 DK020541, R-01 DK116873, R-01 DK125285, R-01 DK105015, R-01 DK113079, R-01 DK126450, and an Einstein-Sinai DRC Pilot and Feasibility Grant (to GL). We thank Luis Santos for his help with the initial technical steps of the snRNA-seq technique.

## Data Availability

All data are available on request from the corresponding author.

---

**Supplemental Table 1.** Information on the human islet preparations used in these studies.

| Donor Code | Source | Age | Race/sex | Height | Weight | BMI | HbA1c | COD | Experiment |
| --- | --- | --- | --- | --- | --- | --- | --- | --- | --- |
| HP20240-01 | Prodo | 42 | Caucasian Male | 65" | 141 lbs | 23.5 | 5.4% | Stroke | In Vitro |
| HP20245-01 | Prodo | 24 | Hispanic Male | 70" | 140 lbs | 20.0 | 5.4% | Head Trauma | In Vitro |
| HP21189-01 | Prodo | 26 | Hispanic Male | 72" | 197 lbs | 26.3 | 5.4% | Head Trauma | In Vitro |
| HP20234-01 | Prodo | 58 | Caucasian Female | 66 | 166 lbs | 26.9 | 5.5% | Stroke | Islet Graft |
| HP21055-01 | Prodo | 36 | Caucasian Female | 57 | 145 lbs | 31.6 | 5.6% | Stroke | Islet Graft |
| HP21203-01 | Prodo | 25 | Hispanic Male | 70" | 187 lbs | 26.5 | 5.8% | Head Trauma | Islet Graft |
| HP21197-01 | Prodo | 53 | Caucasian Male | 67" | 206 lbs | 32.4 | 5.5 % | Anoxic Event | Islet Graft |

\*COD = Cause of death

**Supplemental Table 2.** Information on cell Ranger processed data prior to quality control.

| Donor code | Processing | Number of Cells/Nuclei | Mean Reads per Cell/Nucleus | Median Genes per Cell/Nucleus |
| --- | --- | --- | --- | --- |
| HP20240-01 | Single cell | 8,433 | 19,856 | 1,939 |
| HP20240-01 | Single nuclear | 6,904 | 27,307 | 2,052 |
| HP20245-01 | Single cell | 5,535 | 53,915 | 2,574 |
| HP20245-01 | Single nuclear | 3,927 | 97,308 | 1,150 |
| HP21189-01 | Single Cell | 8,635 | 42,977 | 2,399 |
| HP21189-01 | Single nuclear | 1,683 | 128,235 | 1,972 |
| HP20234-01 | Islet Graft SN | 4,500 | 79,254 | 1,848 |
| HP21055-01 | Islet Graft SN | 2,866 | 106,501 | 1,537 |
| HP21203-01 | Islet Graft SN | 4,930 | 69,792 | 2,601 |
| HP21197-01 | Islet Graft SN | 909 | 349,619 | 2,945 |

\*SN = Single nuclear

